# u-track 3D: measuring and interrogating dense particle dynamics in three dimensions

**DOI:** 10.1101/2020.11.30.404814

**Authors:** Philippe Roudot, Wesley R. Legant, Qiongjing Zou, Kevin M. Dean, Tadamoto Isogai, Erik S. Welf, Ana F. David, Daniel W. Gerlich, Reto Fiolka, Eric Betzig, Gaudenz Danuser

**Affiliations:** Lyda Hill Department of Bioinformatics, UT Southwestern Medical Center. Dallas, Texas, U.S.A.; Department of Biomedical Engineering, University of North Carolina and North Carolina State University, Chapel Hill and Raleigh, NC, U.S.A.; Department of Pharmacology, University of North Carolina, Chapel Hill, NC, U.S.A.; Aix Marseille University, CNRS, Centrale Marseille, I2M, Turing Centre for Living Systems, Marseille, France; Institute of Molecular Biotechnology of the Austrian Academy of Sciences. Vienna, Austria; Department of Molecular & Cell Biology, University of California. Berkeley, California, U.S.A.

## Abstract

Particle tracking is a ubiquitous task in the study of dynamic molecular and cellular processes through microscopy. Light-sheet microscopy has opened a path to acquiring complete cell volumes for investigation in 3-dimensions (3D). However, hypothesis formulation and quantitative analysis have remained difficult due to fundamental challenges in the visualization and the verification of large and dense sets of 3D particle trajectories. Here we describe u-track 3D, a software package that addresses these two challenges. Building on the established framework of particle association in space and time implemented for 2D time-lapse sequences, we first report a complete and versatile pipeline for particle tracking in 3D. We then present the concept of dynamic region of interest (dynROI), which allows an experimenter to interact with dynamic 3D processes in 2D views amenable to visual inspection. Third, we present an estimator of trackability which defines a score for every trajectory, thereby overcoming the challenges of trajectory validation by visual inspection. With these combined strategies, u-track 3D provides a framework for the unbiased study of molecular processes in complex volumetric sequences.

## Introduction

Light-sheet fluorescence microscopy (LSFM) achieves three-dimensional (3D) imaging with minimal phototoxicity, fast sampling, and near-isotropic resolution^1,2^. Given these advances, many dynamic intracellular processes that were once challenging to study even in 2D (e.g., mitosis, cytoskeleton organization, vesicle trafficking, molecular interactions), can now be monitored in the entire cellular volume^1–4^. While the application of computer vision techniques are well established for interrogating cell biological processes in 2D microscopy^5^, these tools do not provide a solution toward the interpretation of dense arrangements of structures moving in a dimensionally unconstrained manner. In 3D, both the visualization of measurement results and their validation require a new set of computational tools. Indeed, a key challenge for image analysis in 3D is the limited ability for a user to interact with the data. The manipulation of time-lapse 3D image volumes is often cumbersome, and any of the projection mechanisms necessary to map the 3D volume into a 2D data representation on a screen is prone with artifacts that may cause erroneous conclusions^6^. Thus, computational tools for 3D image analysis must be able to reveal the complexity of 3D cellular and sub-cellular processes, while being as automated as possible to avoid selection and perception biases.

The most elementary way to measure the behavior of intracellular processes in space and time is particle tracking. Particles can comprise sub-diffraction sized objects that appear in the image volume as bona fide spots, objects of an extended size that appear as a rigid structure, and larger deformable objects. The more complex the object’s shape is, the more sophisticated methods are needed for particle detection. The problem of particle tracking is then defined as the reconstruction of a set of trajectories across time points given the coordinates [x(t), y(t), z(t)] of the identified particles in the individual time points. Many approaches have been proposed to solve this problem ^7–10^. However, only a few of these methods have been implemented in 3D and none of those approaches tackle the visualization and interpretation challenges ^7,11–13^.

Building upon our previous particle tracking work^2,9,14^, we designed the software package u-track 3D to enable the measurement, observation and validation of dynamic processes in 3D (Fig. 1). u-track 3D can detect and track morphologically and dynamically diverse cellular structures, including single molecules, adhesion complexes, and larger macromolecular structures such as +TIP protein complexes associated with growing microtubules. The software design is open, allowing users to import the coordinate files from other detection routines and then apply the u-track framework only for trajectory reconstruction. To overcome challenges inherent to volumetric data, we introduce an extensive library for visualization and mapping of dynamic region of interests (dynROIs) that move with the biological structure under evaluation and enable an intuitive visualization of particle behaviors. Finally, as it is generally impossible to visually validate the trajectory reconstruction in 3D, we present an algorithm for automatic assessment of particle *trackability* throughout the image volume.

**Figure 1:**
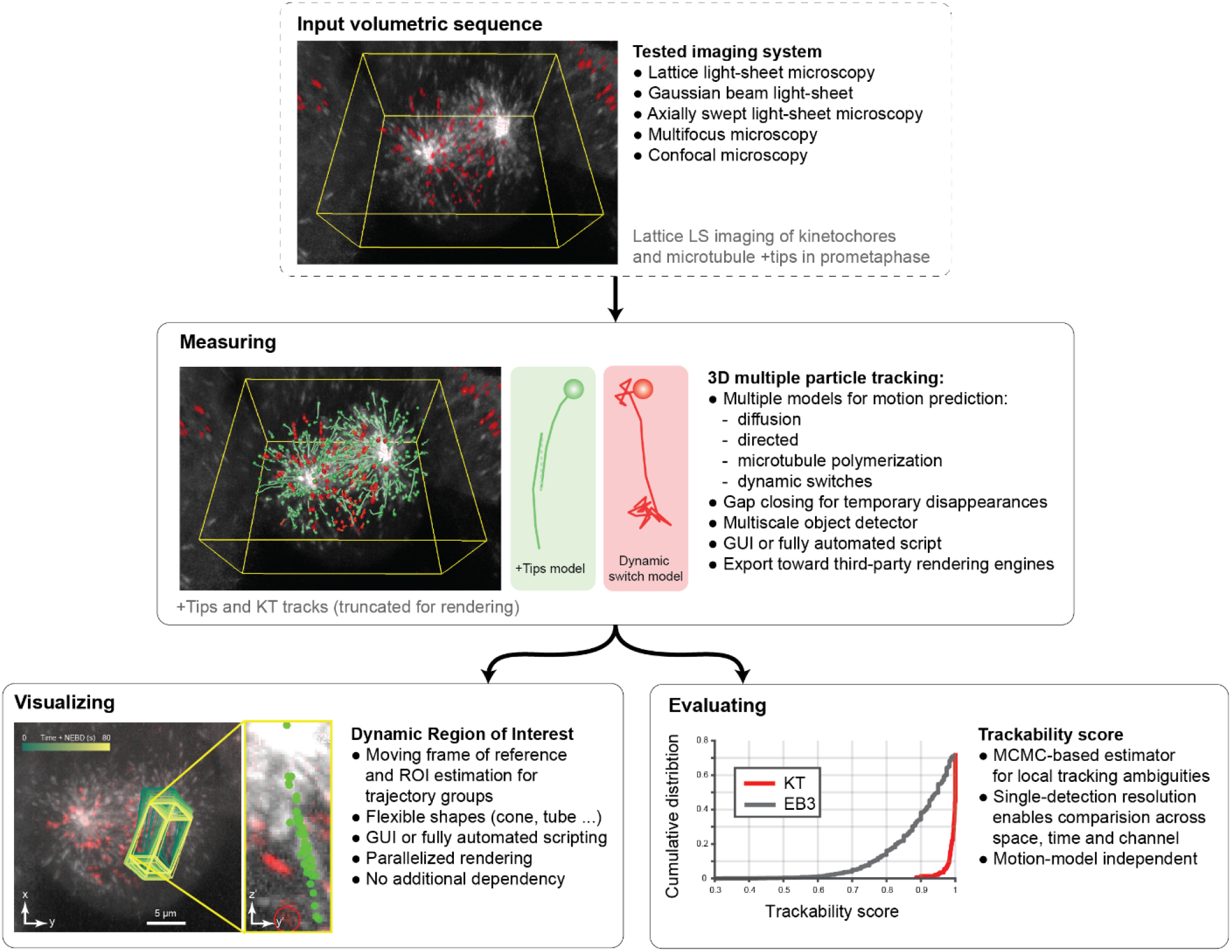
u-track 3D is a complete pipeline for the measurement, visualization, and evaluation of large sets of 3D trajectories. The pipeline is illustrated on lattice light-sheet imaging of HeLa cells undergoing mitosis labeled with eGFP-labeled EB3 and mCherry-labeled CENPA.

## Results

### The u-track 3D pipeline is designed to measure, visualize and validate complex entanglements of 3D trajectories

#### Multiple particle tracking

To generate a robust 3D particle tracking package, we adopted and modified features that were critical for accurate particle tracking in 2D^9^. This includes the break-down of trajectory reconstruction into a frame-by-frame association of corresponding particles followed by an association of the resulting track segments into full-length trajectories. Both steps rely on the same solution for optimal one-to-one assignments of particle detections and track segments in a bipartite graph^9,15^. The two-step approach permits the closing of temporal gaps in the detection of a particle, as well as the handling of particle merging and splitting events that are inherent to many biological processes. As such, the resulting set of trajectories is a global optimum in space and time for a given set of detections. Moreover, u-track 3D incorporates a Kalman filtering approach to model on the fly the characteristics of a particle’s Brownian, directed, and heterogeneous motion, which supports both the procedure for frame-by-frame particle association and the one of track segment association. To support the concurrent tracking of objects of variable sizes we implemented a multiscale particle detector equipped with a generalized adaptive thresholding approach (see Section “Multiscale particle detector” in Material and Methods).

### Dynamic regions of interest

Moving from 2D to 3D images complicates the interaction of a human observer with both raw and derived data. Widely used global image projections, including maximum intensity projection, and other volume rendering techniques are limited by the overlap of many dynamic structures along the viewing axis^6^. However, detailed visualization of 3D images and trajectories in their local context is essential for a user to adjust software control parameters and to interpret the underlying biology. As such, projection approaches have to be tailored to emphasize a subset of selected voxel or aspects of highest interest. Such projections should not only bring the particle or group of particles of interest into focus, but also continuously adapt as the particles move. To meet this requirement, u-track 3D incorporates a framework for rendering particle-centric dynamic regions of interest (dynROIs), thereby allowing the user to follow the particle behavior throughout its lifetime in a visually-comprehensible format. DynROIs are implemented in a hierarchical object structure across molecular, macromolecular and cellular scales (see Section “Dynamic Region of Interest estimation” in Material and Methods). First, u-track 3D provides a variety of shapes (rectangle cuboids, spheres, cones, tubes and rounded tubes) to define region of interest made of one, two or three trajectories. Second, to manage larger sets of tracks, dynROIs are built by estimating an affine transform between the associated point cloud in consecutive time points. Finally, the top level dynROI is defined for the cell. For example, cells embedded in a 3D environment are often randomly oriented, and their orientation changes throughout time. While image-based registration can be used to correct changes in cell orientation, it is computationally expensive, especially as the size of the volume and length of the sequence grows. To reduce the computational burden, we segment and transform the cell mask into a randomly down-sampled point cloud, which is then used to estimate an affine transform.

#### Trackability Score

Validation of tracking results is crucial for proper parameter adjustment during image acquisition and analysis as well as the biological interpretation of integrated measurements. However, it remains an extremely challenging task in 3D datasets, particularly when the particle density is high. Contrary to a scenario in 2D where a single field of view presents a wide range of trajectories for visual inspection, dynROIs in 3D tend to capture only a few trajectories and cannot represent the heterogeneity of local image quality, particle density and dynamic properties, which all affect the tracking accuracy. To solve this problem, we complement u-track 3D with the computation of a local trackability score. For a scenario of homogeneous particle density and directionally random displacements, we offered in previous work a model to compute the probability of tracking errors^2^. However, in a more realistic model, each trajectory bears its own level of uncertainty based on its own stochastic footprint and the configuration of neighboring particles. Here, we compute for every trajectory and every time point the confidence by which the algorithm was able to assign the chosen particle to the trajectory (see Section “Stochastic programming for the evaluation of trackability” in Material and Methods). Specifically, we exploit the particle history, the detection accuracy and the associated motion model(s) to derive a trackability metric that represents the likelihood of each of the chosen associations vis-à-vis the set of alternative associations with neighboring particles. We demonstrate the performance of the resulting score and how it can be used to compare trackability across space, time and the molecules under study.

### 3D MPT combined with light-sheet microscopy simplifies the measurement of endocytic lifetime but gap closing remains a crucial step

To assess the performance of u-track 3D, we investigated the dynamics of various cellular structures imaged by light-sheet microscopy (Figure 2). As reported with u-track, gap closing is a crucial step in 2D particle tracking because of the frequent, transient disappearances: particles might not be detected, particles move in and out of the microscope’s in-focus volume, or particles can temporarily overlap in space. While the latter two sources of disappearance are largely eliminated by proper 3D imaging, the challenges of false or missing detections remain. To test the performance of u-track 3D in closing gaps, we examined the lifetimes of clathrin-coated structures forming at the cell plasma membrane (Fig 2a-c). These structures represent mostly sub-diffraction objects, i.e. they appear in an imaging volume as 3D point-spread functions. We used high-resolution diagonally swept light-sheet microscopy^2^ to sample every second a full volume of puncta generated by the GFP-labeled AP2 subunit of the endocytic coat. In this volumetric sequence, u-track 3D recovered the canonical lifetime distributions of abortive and maturing clathrin-coated pits, that is, an exponential decay for abortive pits and Rayleigh-like distribution with maximal probability around 20 s for maturing pits (Fig 2.c, Movie 1 and Section “Clathrin-mediated endocytosis study on a glass coverslip” in Materials and Methods). While the identification of those two populations has required extensive trajectory analysis to discount incomplete trajectories caused by the limitations of 2D microscopy (e.g. the transient arrival of golgi-associated clathrin coated structures into the evanescent illumination field)^16^, our u-track software achieves accurate trajectory classification directly by thresholding the maximum intensity of trajectories in 3D (Fig 2.b,c). Importantly, the separation of lifetime distributions into two pit classes can only be obtained with the support of gap closing (Fig 2.c), suggesting that gaps present a hurdle for accurate tracking also in 3D.

**Figure 2:**
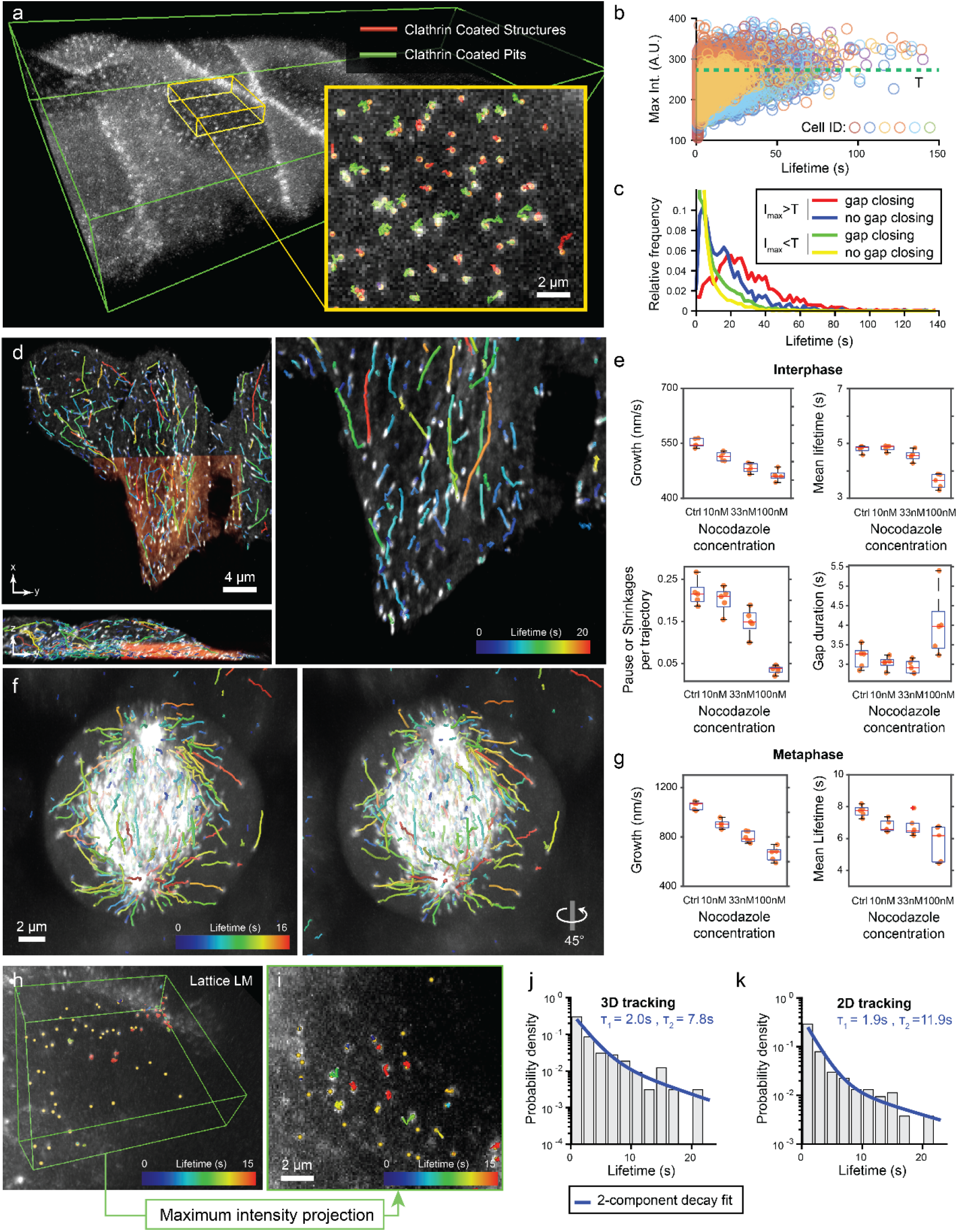
u-track 3D supports a variety of imaging and biological scenarios. a) Maximum intensity projections (MIP) of a rat kidney cell layer imaged with diagonally scanned light-sheet microscopy (diaSLM). Cells are expressing eGFP-labelled alpha subunit of the AP-2 complex. Green box is 160×40×12 um. Inset shows trajectories of clathrin aggregates classified as clathrin coated structures or maturing pits. b) Normalized maximum intensity of each trajectory as a function of lifetime plotted for six cellular layers composed of multiple cells each. The green line denotes the median of the cumulated distribution (value T). c) Probability density of lifetime for the set of trajectories above and below the threshold value T, with and without gap closing (N=6 cellular layers, pooled trajectories lifetimes). d) Maximum intensity projection (MIP) of HeLa cells in interphase imaged with lattice light-sheet microscopy (LLSM) expressing eGFP-labeled EB1 (orange area is 30×32×7 um). Overlay highlights EB1 trajectories. e) Average microtubule lifetimes, microtubule growth rate as well as average number and duration of pause and shrinkage events per trajectory for increasing concentrations of nocodazole (N = 5 per conditions; center line, median; box limits, 25 and 75 percentiles; whiskers, extremum). f) MIP of HeLa cells in metaphase imaged with LLSM along with 45-degree rotation around the vertical axis. Overlay highlights EB1 trajectories. g) Same as e) measured for cells in metaphase (N = 5 per conditions). h) MIP of mouse embryonic stem cell (ES) nucleus imaged with LLSM expressing GFP-labeled transcription factors. Green box is 13×13×3 um. Overlay highlights SOX2 trajectories. i) MIP of ES cell nucleus imaged with LLSM expressing GFP-labeled transcription factors. Overlay highlights SOX2 trajectories tracked after MIP transformation. j) Probability density of SOX2 binding time measured in LLSM overlaid with a 2-component decay fit (N=1 cell). k) Probability density of SOX2 binding time measured in projected LLSM data overlaid with a 2-component decay fit (N=1 cell).

### Directed motion filtering captures drug-induced variations in microtubule polymerization in dense 3D spindle imaged with lattice light-sheet microscopy

With limited sampling frequency in volumetric imaging due to the serial acquisition of a z-stack comprising tens to hundreds of focal slices, the automated reconstruction of particle trajectories can be improved by dynamic motion models through Kalman filtering. To assess the performance of a 3D implementation of previously published motion models for 2D tracking of microtubule polymerization dynamics^17,18^, we imaged and tracked the dynamics of microtubules in HeLa cells by following GFP-fusions of the microtubule plus-end tracking protein EB1 using lattice light-sheet imaging^1^ at 1 Hz in interphase and metaphase. We quantified metrics such as growth rate, growth lifetime and pause frequency (see Section “Microtubule instability measurement” in Material and Methods). The latter is a measure for the probability that a stalled or shrinking microtubule, which is accompanied by disappearance of the EB1 particle in the movie, is rescued to renewed growth (see Supplementary Figure 1 and Movie 2). Consistent with our previous observations in 2D^17^, u-track 3D faithfully detected a dose-dependent decrease in all three metrics upon treatment of cells with the microtubule-destabilizing drug nocodazole (Fig 2.d-e). We also investigated the destabilizing effect of nocodazole by measuring the number and duration of pauses or shrinkages that occur with drug treatment (Fig 2.e) until disappearance at the highest concentration. We then extended our analyses to mitotic cells, where the density of EB1 particles is much higher in central regions of the mitotic spindle (see Movie 3). Both scenarios show a strong response in nocodazole concentration, indicating that u-track 3D properly captures the drug-induced variation of growth rate and lifetime (Fig 2.f-g), despite strong variations in particle density.

### 2D measurements artificially increases the lifetime of interaction between Transcription Factor and chromatin

We then sought to investigate the impact of the depth information on the measurement of biological quantities when compared to 2D particle tracking. We compared the u-track gap closing and motion modeling capacities, including an approach to follow particle trajectories with erratic go-stop-go behaviors^14^, in both 2D and 3D data using single molecule imaging. We used a lattice light-sheet microscope to image the interactions between transcription factors (TFs) and chromatin in embryonic stem cells. In a study using the same biological system, Chen *et al*^19^ had shown that TFs alternate between short-lived binding events at non-specific chromatin sites (residence time ∼0.75s), 3D diffusion (average duration ∼3s) and longer lived transcription events where the TF is bound at specific chromatin sites (residence time ∼12s). These results were quantified in 2D using both light-sheet and widefield imaging. We performed the same analysis, now applying 3D tracking, and contrasted the results to the tracking of 2D projections of the same 3D volumes (Fig 2.h-k and Section “Single molecule dynamics study with lattice light-sheet microscopy” in Materials and Methods). Interestingly, when analyzed in 3D, the residence time of binding events was reduced by one third (∼7.8s in 3D vs 11.9s in 2D). Thus, only on 2D projections are we able to reproduce the results of the original study, which differ significantly from the 3D results. Interestingly, the shorter binding time observed in 3D trajectories is consistent with measurements performed in nuclear receptors studies^20,21^. Together, these data suggest that the overlap caused by axial integration of the fluorescent signal imaged in 2D may artificially prolong the lifetimes and change the conclusion on binding kinetics.

### 3D adhesion formation visualization and analysis using dynamic region of interest suggests that rounder adhesions are closer to collagen fibers

We illustrate the application of a whole-cell dynROI with the study of spatial interactions between cell-matrix adhesions and fluorescently labeled 3D collagen fibers in osteosarcoma cells imaged by axially swept light-sheet microscopy^3^ (Fig 3.a and Movie 4). The dynROI allowed us to visualize the relationship between adhesion shapes and its proximity to collagen fibrils, showing two populations of globular and elongated adhesions (Fig 3.b-d). The most elongated adhesions are located predominantly at the tip of the pseudopodial extensions and align with the protrusive direction, while the round adhesions concentrate in the quiescent part of the membrane. Our measurements show that this elongation distribution can be decomposed further (Fig 3e). We found a unimodal distribution of mostly globular adhesions in close contact with collagen fibers, assessed by a score that integrates the distances between voxels in adhesions and collagen fiber masks (see Section “Adhesions and collagen interaction imaging and analysis” in Materials and Methods). In contrast, adhesions with a lesser degree of collagen contact display a bimodal distribution of globular and elongated adhesions. These data suggest – quite unexpectedly from what is known in 2D – that the most engaged adhesions may be the least elongated^22^. We further conjecture from this finding that adhesion elongation in 3D may be less driven by a zippering of an integrin-mediated plaque along a collagen fiber, but rather dictated by the organization of cell-cortical actin fibers or the local collagen architecture. Indeed, this behavior becomes apparent by replay of time-lapse sequences of the proximity and elongation parameters in the spatially stabilized dynROI (Movies 5, 6 and 7). DynROIs are thus a powerful way to assess the spatial distribution and heterogeneity of molecular interactions in highly dynamic cells.

**Figure 3:**
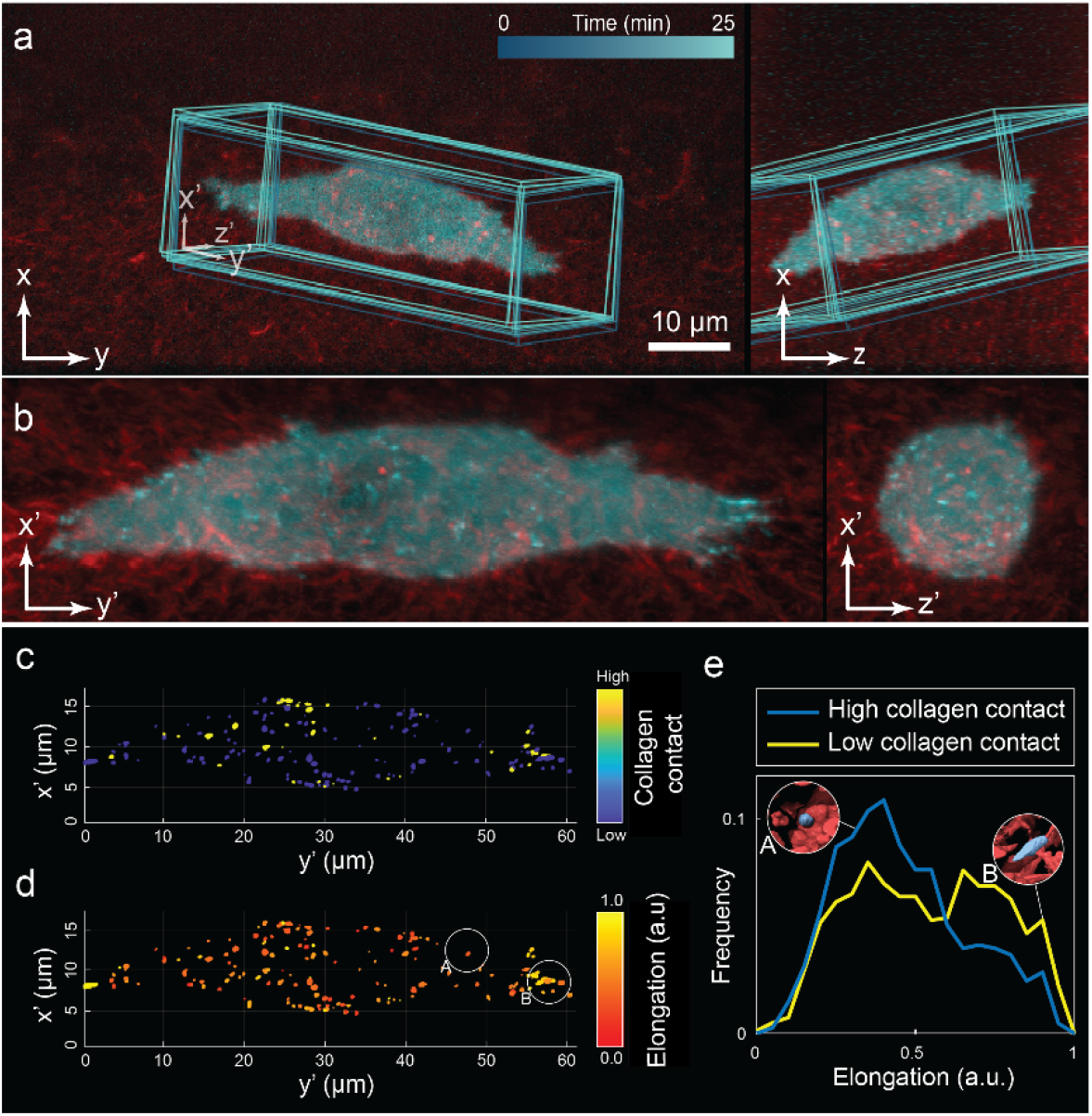
Dynamic Regions of Interest (dynROI) reveals the behavior of local particle involved in highly dynamic sub-cellular processes. a) Dual-colored orthogonal MIP of osteocarcinoma cells expressing eGFP-labeled paxillin and embedded in collagen labelled with Alexa fluor 568. Overlay highlights dynamic region of interest (dynROI). b) View of the dynROI. c) – d) Detection of adhesions colored as a function of the degree of collagen contact and elongation. e) Probability density of elongation for adhesions with high and low degree of contact with collagen fibers (N=1 cell).

### Dynamic region of interest enables the automated exploration of mitotic spindle across scale and the observation of different kinetochore capture mechanisms by microtubules

Many cellular processes involve a massive reorganization of multiple macromolecular structures, which challenges 3D analysis by conventional visualization approaches. A classic example is the mitotic spindle of vertebrate cells^23^. While thousands of microtubules reorganize to form a dense bipolar array, the two spindle poles move apart and rotate back and forth. At the same time, spindle microtubules establish contacts with chromosomes at specialized attachment sites termed kinetochores, and subsequently move chromosomes towards poles or the spindle center. The complexity of these structures and their rapid dynamics are virtually impossible to understand by mere visual inspection of volume renderings. We therefore assessed how u-track 3D and dynROIs facilitates the analysis of this process. The image dataset comprises dual-channel time-lapse sequences of GFP-labelled microtubule plus-ends and mCherry-labelled kinetochores of mitotic HeLa cells, acquired by lattice light sheet microscopy as described in ref^24^, with an acquisition frequency of 0.1 Hz to enable longer imaging, from prometaphase to metaphase. In addition to microtubule plus-ends and kinetochores, our multiscale particle detection approach is able to locate the spindle poles based on the diffuse clouds of microtubule plus-end marker. Pole trajectories can then be used to define a dynROI that follows the spindle motion (Fig 4.a and Section “Dynamic Region of Interest estimation” in Materials and Methods). An embedded second dynROI follows the point cloud formed by the kinetochore trajectories (Fig 4.b and Section “Dynamic Region of Interest estimation” in Materials and Methods). Based on the pair of dynROIs, we further construct a planar dynROI with an orientation that is defined by the interpolar axis and a vector following the kinetochore-associated dynROI motion (Fig 4.c,d, Movie 8, 9 and Section “Dynamic Region of Interest estimation” in Materials and Methods). Our framework for dynROI estimation thus enables the automated visualization of dynamic mesoscale structures composed of different molecular assemblies.

**Figure 4:**
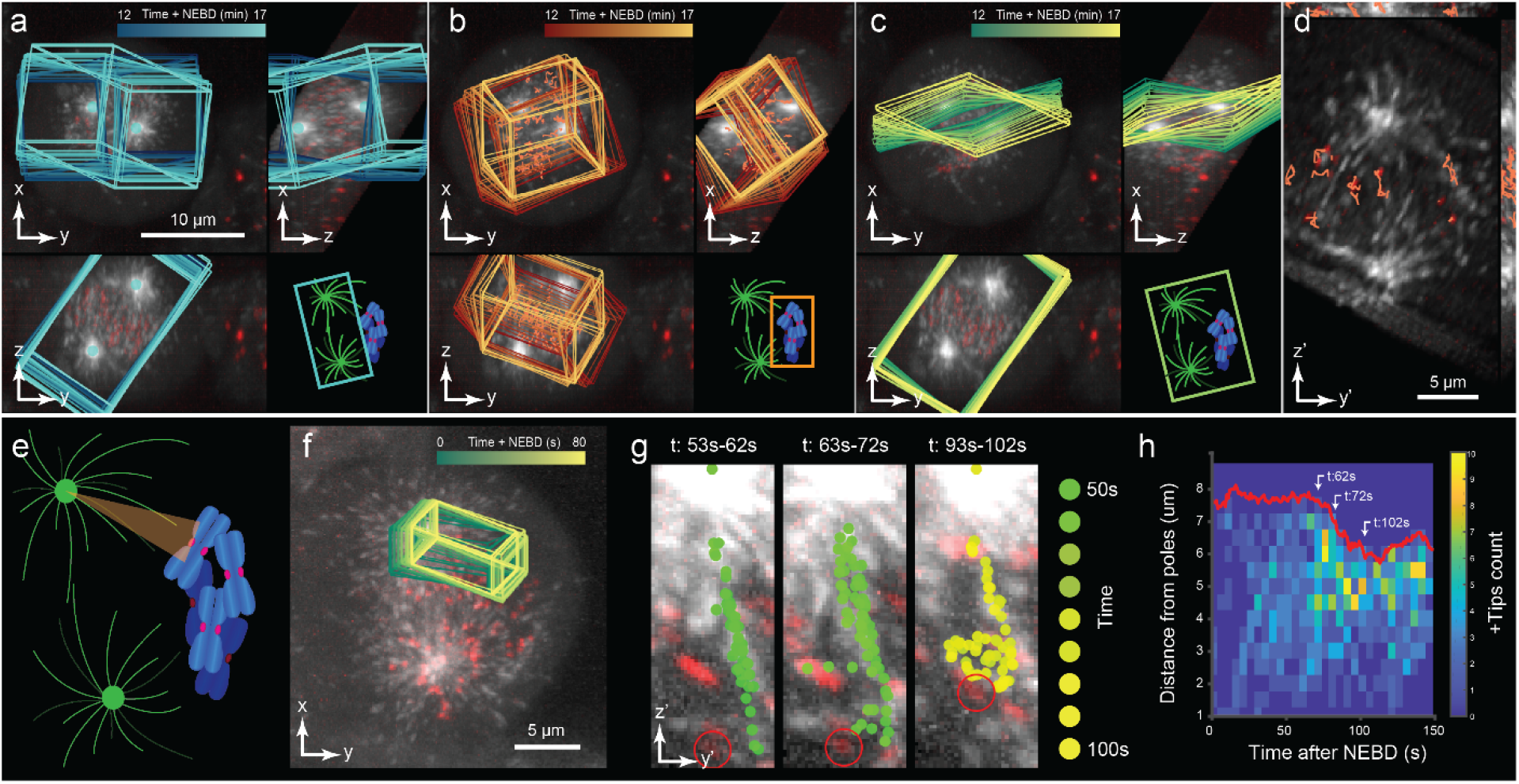
a) - d) Dual-colored orthogonal MIP of HeLa cells undergoing mitosis labeled with eGFP-labeled EB3 and mCherry-labeled CENPA. Overlays highlight, a) a dynROI built around centrosome trajectories, b) a dynROI built around CENPA trajectories, and c) a plane built to visualize the dynamics of chromosomes relative to the spindle location. d) View of the dynROI following described in h. e) Definition of a conical dynROI between a centrosome and a kinetochore. f) Dual-colored orthogonal MIP of HeLa cells during pro-metaphase. Overlay highlights the motion of the dynROI. g) Cumulative overlays of the detected microtubule plus-end position for three periods of 10 seconds between 53s to 102s post nucleus envelope breakage. h) Plus-ends count function of time and distance from the pole (N = 1 dynROI).

The hierarchical model of dynROIs also enables the detailed analysis of microscale collective molecular processes that move throughout the cellular volume. This facilitates the study of complex subprocesses, as for example the formation of kinetochore fibers during spindle assembly. In previous work^24^, we showed with spindle-wide statistics and indirect modeling that kinetochore fiber formation is accelerated by an augmin-dependent nucleation and directional growth along the fiber towards kinetochores. To expand the analysis of this process, we now use dynROIs to directly visualize the dynamic space between spindle poles and kinetochores (Fig 4.e-g and Movie 10). We define a kinetochore fiber assembly dynROI by a cone whose medial axis connects spindle pole and a target kinetochore (see Section “Dynamic Region of Interest estimation” in Materials and Methods). By systematically inspecting kinetochore fiber assembly dynROIs, we observed a strong directional bias in microtubule polymerization along a kinetochore fiber and toward kinetochores, consistent with previous observations^24^. We also observed microtubule polymerization branching off from a kinetochore fiber and polymerizing toward another kinetochore (circled in red in Fig 4.g, time 53 s – 72 s). The branching was followed by a rapid poleward motion of the targeted kinetochore and an increase of plus-end count in the dynROI (Fig 4.g,h, time 93 s – 102 s) suggesting that the target kinetochore was captured, and that this new capture established a new avenue for microtubule amplification. This data indicates that kinetochore capture might occasionally involve interactions between neighboring kinetochore fibers. The example underscores how the dynROI library implemented in u-track 3D enables the visual discovery of dynamic processes that are completely obscured in 3D image volumes.

### The trackability score detects tracking ambiguities with high precision

Given how cumbersome it is to visualize particle trajectories, a systematic validation of the tracking performance by visual inspection – the gold standard in most cell biological studies in 2D – seems out of reach for many applications in 3D. Thus, we developed a self-assessment pipeline that attributes every trajectory with a trackability score predicting the accuracy of the automated tracking solution. The principle behind the trackability score and its performances are explained in Figure 5 and Section “Stochastic programming for the evaluation of trackability” of Material and Methods. Figure 5.a shows an example of an ambiguous association between time *t* − 1 and *t* where two assignment hypotheses between detections and track heads have a similar association cost. Despite this ambiguity, the bipartite graph matching identifies a single optimal solution without the possibility for error estimation (see Fig 5.a.i). To determine the level of ambiguity, we resample all track head predictions *N* times and test the stability of the original assignment (one resampling example is shown in Fig 5.a.ii). The approach is illustrated in Figure 5.b-d based on the tracking of a transcription factor in a dataset acquired with multifocus microscopy and previously published in ref^25^. Each dot indicates a resampled prediction of the particle location at *t*, and blue versus red defines whether the newly computed local assignment matches or differs from the original solution. The trackability score is defined as the fraction of matching samples. As such, the trackability score infers tracking accuracy by considering both the local competition of detection candidates for track head associations and the uncertainty of motion prediction for each track head.

**Figure 5:**
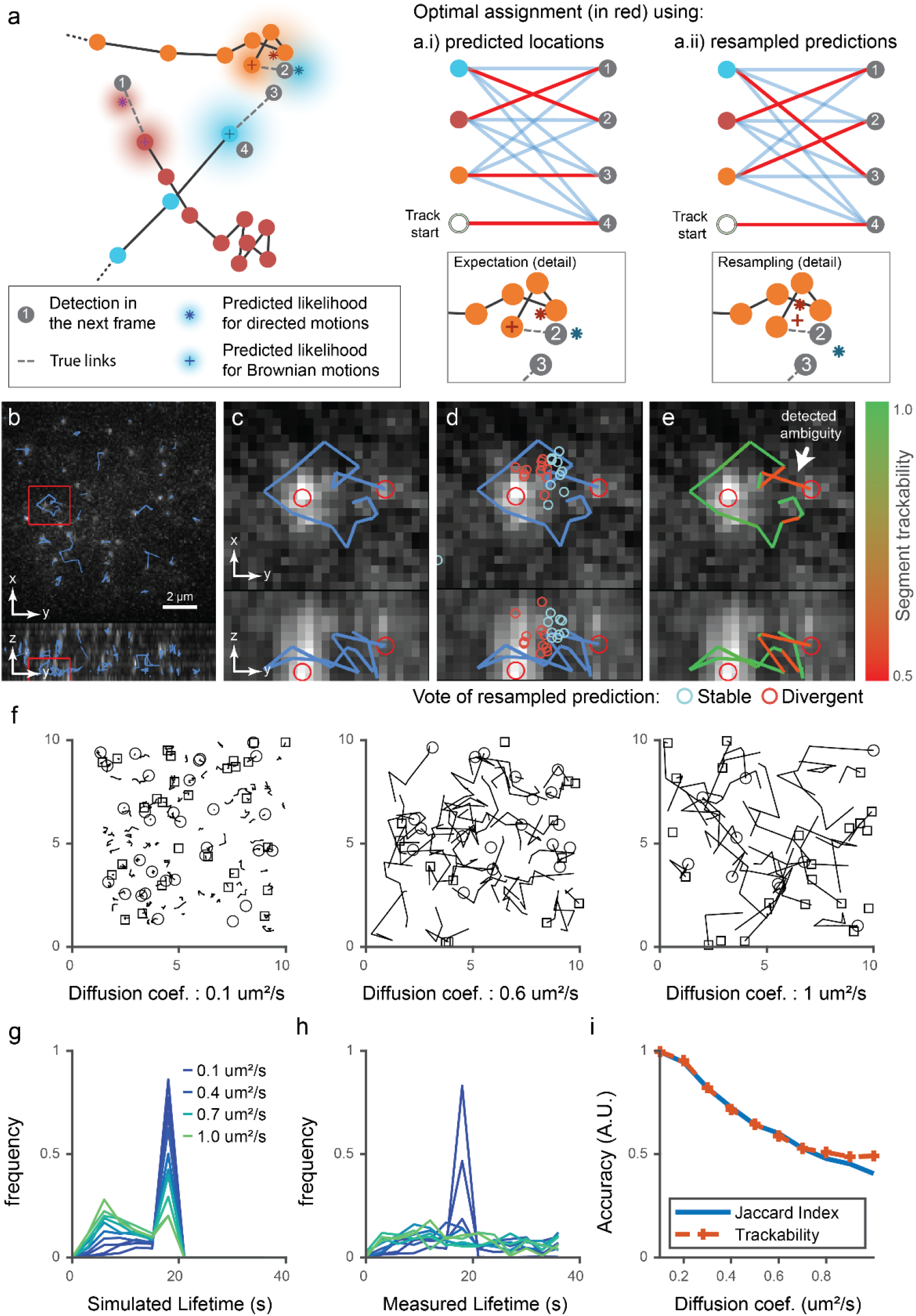
The trackability score relies on the stochastic footprint of each trajectory to infer tracking accuracy. a) Example of a tracking ambiguity due to three trajectories in close proximity (orange, blue and red). Dashed lines represent the true motion between track heads at time *t* − 1 and detections at time *t*, represented by gray dots. Colored gradients represent the likelihood of each expected particle location at time *t*, estimated using the history of positions up to time *t* − 1 and considering multiple motion model hypotheses. The optimal assignment between the expected and detected particle positions at time *t* in this case yields an erroneous assignment from the orange track head to detection 2 and from the blue track head to detection 3 (graph a.i). Resampling of the expected locations results in a new assignment (graph a.ii), this time without error. b) Orthogonal Maximum Intensity Projection (MIP) of Embryonic Stem (ES) cells expressing eGFP-labelled Sox2 molecules imaged by Multifocus Microscopy. Overlaid boxes highlight the ROI enlarged in c – e). c) Orthogonal MIP of ROI. Overlay shows a trajectory where two close detections create assignment ambiguity. d) Overlay illustrates the stochastic resampling of the predicted particle positions at this time-point; blue circles: assignments in agreement with the original solution; red circles: assignments that differ from the original solution. e) Overlay shows trajectory segments colored according to estimated trackability scores. g) Examples of simulated trajectories with diffusion coefficients ranging from 0.1 um^2^/s to 1 um^2^/s with a fixed particle density of 0.1 um^-3^. Visualization is limited to five consecutive frames to reduce clutter. f) Lifetime of simulated trajectories (the change in distribution is due to trajectories leaving the field of view as the diffusion coefficient increases). h) Lifetime distribution measured through tracking shows a loss of the original distributions when the diffusion coefficient exceeds 0.2 um^2^/s. i) Accuracy measured through the jacquard index (JI, blue); the trackability score (orange, dashed), which is derived without external ground truth, closely follows the JI up to a diffusion coefficient 0.6 um^2^/s beyond which tracking is random.

We evaluated the capacity of our score to predict tracking quality in a variety of scenarios. We simulated multiple sets of trajectories of increasing complexity and applied u-track 3D to trace the particle movements between time points. Using the ground truth, we then classified each link of the extracted traces as a true positive (TP) or false positive (FP). This classification allows us to compute for each simulated sets the true Jaccard Index (JI), a popular metric for linking accuracy^7^, computed as 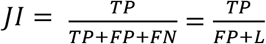 where L represents the number of simulated links. Of note, we did not introduce noise to the set of simulated particle detections to focus our evaluation on the ability to predict the risk of linking errors. Figure 5.f presents an example of simulated trajectories with increasing diffusion coefficients; the display is truncated to five consecutive time points to improve visibility. With a detection density fixed to 0.1 um^-3^ and increasing speed of diffusion, the tracking performance rapidly deteriorates to completely inaccurate links and trajectory lifetime distributions (see Fig 5.g-h). The trackability score follows precisely the decrease in the JI until it plateaus at 0.5 for very challenging conditions (Fig 5.i). The initially close match between trackability score and JI is expected by design as larger diffusion speeds increase ambiguities as well as the number of false positives and negatives.

However, beyond a diffusion of 0.6 um^2^/s, the prediction of the particle location in the next frame is less likely centered on the correct detection. As such, during the resampling of the expected particle location, the rate of samples in agreement versus disagreement with the original link is defined by chance, hence the score plateaus at 0.5. We also simulated a scenario in which the particle density increases at a diffusion fixed to 0.3 um^2^/s. Analogous to the increase in diffusion, the trackability score follows the JI up to a density of 0.25 um^-3^ where the two performance measurements start to diverge (see Supplementary Figure 2). In the case of directed displacements and a given density of 0.1 um^-3^, our trackability score also follows the true JI up until a critical velocity of 1.8 um/s which is more than twice the average distance between a particle and its closest neighbor (see Supplementary Figure 3). Finally, we sought to test our approach in a scenario in which trajectories undergo sudden transitions between diffusive and directed motion (see Supplementary Figure 4). Of note, the densities, diffusion coefficients and velocities are fixed in this scenario and the only parameter that varies is the transition rate, ranging from 0 (no transition) to 0.5 (on average one transition every two frames). Our results show that the trackability score also correctly predicts the reduction in tracking accuracy as increasing transition rates makes tracking more challenging. A quasi-plateau is reached due to the high frequency of dynamic transitions. A detailed description of the parameters used for all simulation experiments can be found in Section “Simulation parameters for trackability evaluation”. In conclusion, our simulations show that the proposed trackability score is able to detect subtle changes in tracking quality in a large variety of scenarios. While our score diverges from the true Jaccard Index when the simulated frame rate is not high enough to initialize motion prediction, the trackability score nevertheless detects at least 60% of linking errors, indicating that the tracking experiment must be re-designed for accurate estimation of both trajectories and errors.

### The trackability score can be used to compare tracking quality across time, space and fluorescent channels

In order to evaluate the performance of the trackability score in indicating high vs low confidence in experimental tracking results, we first analyzed the spatiotemporal variation in the tracking quality of endocytic pits moving along a dynamic membrane (see Section “Endosome trackability on cell cultured on top of collagen” in Material and Methods). The cells were plated on collagen and imaged at high-resolution using diagonally swept light-sheet microscopy^2^. The acquisitions present a large variety of dynamic behaviors, from a quiescent membrane in the center to rapid protrusive displacements at the leading edge (Fig. 6.a). To selectively analyze those areas, we manually selected dynROIs to capture a quiescent area, a slow protrusion-retraction cycle and a fast protrusion-retraction cycle. Since both the cell and its environment are moving, those dynROIs were selected within a larger dynROI built from all the trajectories detected (see Movie 11). The trackability score shows a consistently high score in the quiescent dynROI, a large and transient decrease in trackability around the fast protrusive motion and an average decrease in the slower protrusion (Fig 6.b,c). Our score is thus able to accurately reflect the variation in trackability across space and time and detect time points when tracking ambiguity arises due to rapid movement at the level of the whole cell.

**Figure 6:**
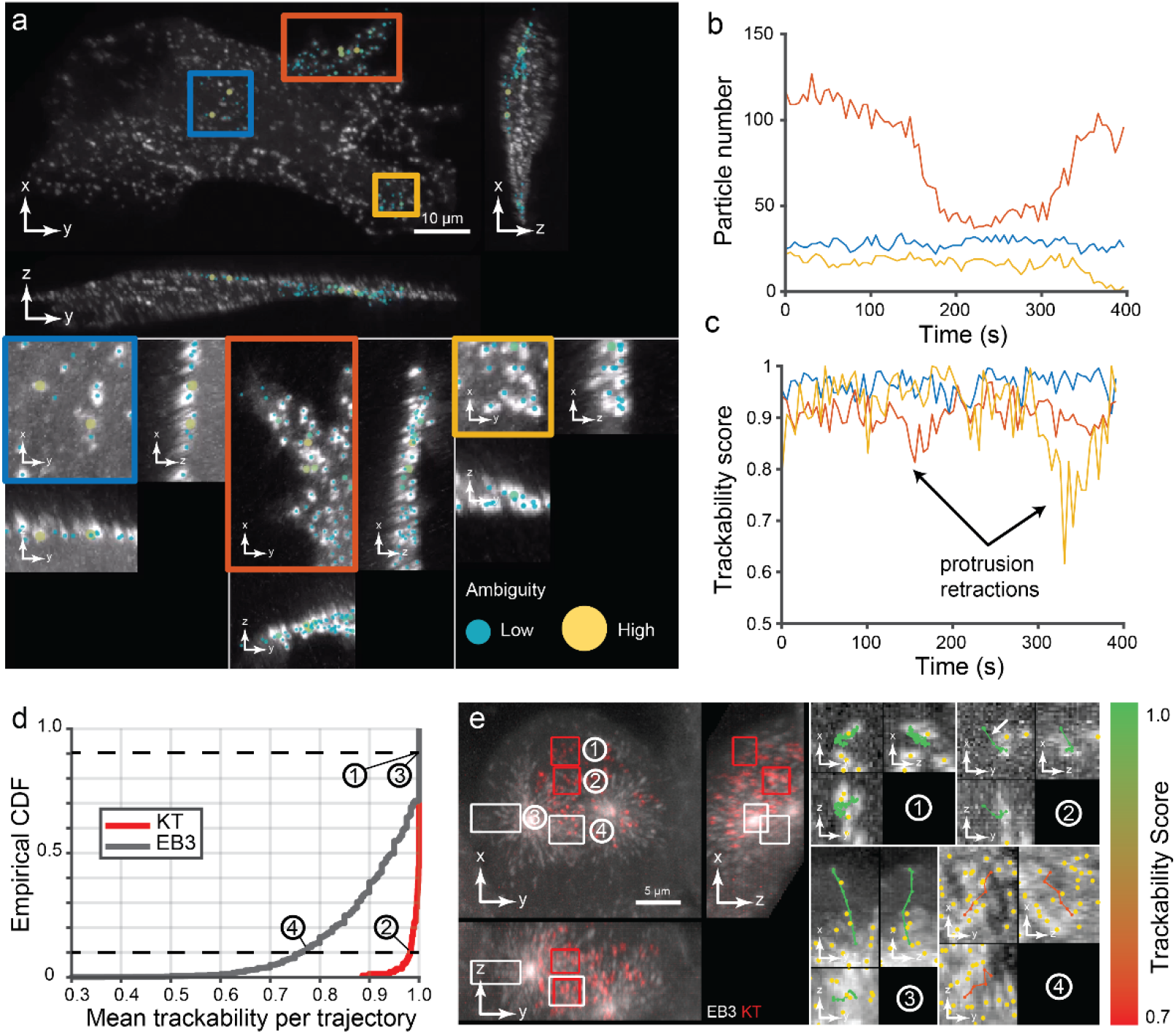
Demonstration of trackability score on experimental data. a) Orthogonal MIP of breast cancer cells imaged with diaSLM expressing eGFP-labelled alpha subunit of the AP-2 complex. Boxes show ROIs with different types of activity. Dot overlays show local level of ambiguity. b) Number of track segments over time for the three ROIs (N=1 cell). c) Trackability score over time for the three ROIs (N=1 cell). d) Cumulative distribution of the average trackability score of trajectories for both EB3 and Kinetochore channels sampling the dynamics of the mitotic spindle shown in Fig 4. e) Four ROIs (two for each channel) showing trajectories colored according to their mean trackability score. Trajectories were selected near the 10th and 90th percentiles of the cumulative distribution. Yellow dots show surrounding detections.

In a second experiment we tested the capacity of our trackability score to compare tracking quality in a complex scene with heterogeneous cellular structures. We analyzed the spindle assembly dataset shown in Fig 4.f-h with labelled microtubule plus-ends and kinetochores. The cumulative distributions of trajectory-averaged trackability scores showed that kinetochore trajectories overall are more reliably reconstructed compared to microtubule plus-end trajectories (Fig 6.d), consistent with their much lower density and slower motion. Our score also enables the analysis of trackability on a per-trajectory basis. Indeed, trajectories with a trackability score near the 90th percentile of the cumulative distribution appear to be error-free for both plus-end and kinetochore channels (Fig. 6.e). In contrast, a plus-end trajectory with a score near the 10^th^ percentile shows a likely erroneous path in an area of very high-density, crisscrossing microtubules. Because of the overall much higher trackability of the kinetochore channel, the 10^th^ percentile of the trackability score distribution of these trajectories picks out a trajectory with only one likely wrong link (see arrow) caused by false positive particle detection. Based on these examples we conclude that the trackability score is a faithful reporter of the overall accuracy of tracking results in a given imaging channel, and it further assists selection of correctly tracked objects in a dense population of trajectories.

## Discussion

We report here a new version of the popular tracking framework u-track, which now enables the study of particle dynamics in 3D live microscopy and tackles key challenges in the exploration and analysis of those complex datasets. First, we demonstrate the 3D implementation of the particle trajectory reconstruction in space and time, including several particle motion models that support particle tracking in a variety of imaging and biological scenarios. We demonstrate applications of the u-track 3D software to the measurement of lifetimes of endocytic events, of microtubule growth dynamics, and of the stop-and-go behavior of individual transcription factors binding to chromatin. We introduce dynROIs, which unveil local particle behaviors embedded in highly dynamic sub-cellular processes that would otherwise be buried in global renderings of the 3D trajectories. We illustrate these functionalities across scales, from the cell-wide heterogeneity of adhesions in a collagen-embedded cell, to the microscale organization of microtubules during chromosome capture. Finally, we introduce a strategy for automatic detection of tracking ambiguities that pinpoints the locations and times of potential tracking errors in the full set of reconstructed trajectories. We demonstrate the approach on simulations and experimental datasets ranging from mapping the spatial heterogeneity of tracking quality in the quantification of endocytic events, to the automatic identification of high- and low-quality trajectories imaged in the mitotic spindle. Thus, u-track 3D complements the development of light-sheet microscopy with a much-needed tool set for the exploration and the quantitative analysis of the biological processes these movies capture.

The u-track 3D software is implemented in Matlab and distributed with a user-friendly GUI and tutorial scripts. The GUI is designed for testing the software and for the interactive visualization of the particle detections, trajectories and dynROI locations overlaid onto the raw data. In particular, both raw voxels and measurements can be observed using either slice-by-slice visualization or maximum intensity projections in the frame of reference of the laboratory or in a frame of reference of a dynamic region of interest. The scripts are primarily used for batch processing and analysis at scale, and they enable the systematic visualization of tracking results across a full dataset with a newly designed renderer. As opposed to traditional interactive rendering, our engine is designed for the fully automated and parallelized rendering of raw data and overlaid measurements that takes advantage of the asynchronous nature of processing jobs. Montages of raw and overlaid images can be easily specified and saved in a variety of formats (png, avi and gifs). As an example, all of the panels in Figure 4 presenting microscopy data have been produced by this rendering engine. The script interface also provides a finer control of the shape of dynROIs than the GUI (cone, tube, rounded tubes etc …). Finally, both detection and tracking can be limited to those dynROIs, enabling the rapid adjustment of algorithm parameters before processing the full dataset and tracking in a more appropriate frame of reference. Two datasets are provided to test the software, one extracted from the endocytosis imaging introduced in Figure 6.a, the other extracted from the mitosis imaging experiment introduced Figure 4.

While the robustness and wide applicability of the software already has been tested in several studies^24,26^, many challenges remain toward a generic approach for the automated exploration of 3D sequences. A chief bottleneck comes with the multiple sources of motions occurring across scales, especially in more physiologically relevant environments with a high degree of freedom. Indeed, while a given framerate may be sufficient to sample and track the motion of particles on a static substrate, the object may not be trackable when the particle-embedding volume moves rapidly. U-track 3D addresses this problem with the estimation of dynamic regions of interest, which allow the pre-alignment of particle groups associated with an entire cell or subcellular structure on a coarser scale for refined tracking of individual particles on a finer scale. However, the automated estimation of the scale, type, clusters and magnitude of those displacements remains an open problem for heterogeneous groups of objects. New developments in stochastic filtering approaches for multiscale displacements are thus necessary. Recent progress in neural networks to mimic Bayesian approaches are promising^27,28^ and could also be adapted to a multiscale representation. Another key challenge in the analysis of dynamic 3D data is the quantification of the motion of diffuse signaling molecules or macromolecular structures that do not present a well-defined particle in the imaged volume. These motions can be estimated coarsely using 3D optic flow approaches for which a few promising methods tailored to fluorescence imaging have been proposed^29–31^. Finally, the visualization and interaction with large multidimensional data remains difficult. In this paper we propose to remove any direct manipulation of the virtual camera to direct the rendering through automatically defined dynROI. While we do believe this is the future of 3D sequence exploration, the underlying rendering engine is limited to maximum intensity projections or slide-by-slide visualization. Community efforts are currently underway to provide a generic and versatile graphic library along with GUI interface such as Napari^32^ and Sciview^33^. They could complete the capabilities of our renderer with more advanced volumetric rendering (alpha, ray casting) as well as surface rendering. We thus introduce u-track as a feature-complete software for the quantification and analysis of 3D trajectories, but also as a steppingstone toward the automated exploration of any types of dynamic datasets. In the meantime, as we deliver the software to the community, we are continuously improving the software by fixing bugs and evaluating suggestions for improvements made by the community.

## Supporting information

Movie description

## Authors contributions

P.R. and G.D. designed the research. P.R. wrote the tracking and rendering software, and performed data analysis. Q.Z. developed the software’s graphical user interface. W.R.L., K.M.D., A.D., T.I. and, E.S.W. performed the biochemical and imaging experiments. K.M.D., R.F. and E.B. provided imaging resources. P.R. and G.D. wrote the manuscript. All authors read and provided feedback on the final manuscript.

## Acknowledgments

The authors are grateful to Yuko Mimori-Kyosue at the RIKEN Institute for the gift of the HeLa cells expressing eGFP-EB1 and fruitful conversations. We are also grateful to Zhe Liu at Janelia Research Campus for giving the E.S. cells. P.R. has been funded by the fellowship LT000954/2015 from the Human Frontiers in Science Program and the « Investissements d’Avenir» French Government program managed by the French National Research Agency (ANR-16-CONV-0001) and from Excellence Initiative of Aix-Marseille University - A*MIDEX.”. Work in the Danuser lab has been funded by the grant R35GM136428. W.R.L. acknowledges support from the Searle Scholars Program, the Beckman Young Investigator Program, an NIH New Innovator Award (DP2GM136653) and the Packard Fellows Program. K.M.D receives partial salary support from R01MH120131, R34NS121873, and R01DK127589. Research in the laboratory of D.W.G. has been supported by the Vienna Science and Technology Fund (WWTF; project nr. LS14-009), and by the Austrian Science Fund (FWF special research program SFB “Chromosome Dynamics”; project nr. SFB F34-06). R.F. has been supported by the Cancer Prevention Research Institute of Texas (RR160057), and the NIH (R33CA235254 and R35GM133522). Lattice light-sheet imaging of the mitotic spindle data produced in collaboration with the Advanced Imaging Center, a facility jointly supported by the Gordon and Betty Moore Foundation and Howard Hughes Medical Institute at the Janelia Research Campus.

## Data and code availability

The latest version of the software described here, a user’s guide for both GUI and scripts and test datasets are available from https://github.com/DanuserLab/u-track3D. The data that support the findings of this study are available from the corresponding authors upon reasonable request.

## Materials and methods

### Multiscale particle detector

Three-dimensional microscopy imposes specific constrains to the design of a particle detector. First, the diversity of shapes and sizes of intracellular structures may not be visible to the naked eye in a volumetric rendering, we must thus design a detector that is responsive to those variations. Second, light scattering and variation in signal intensity create large changes in signal-to-noise ratio (SNR) across space that are also difficult to assess visually. Our detector must then be adapted to those changes from low to high SNR. Finally, the large dimension of 3D data sets requires the design of computationally efficient approaches. Following, we describe a multiscale detector equipped with an adaptive thresholding approach that tests multiple possible scales at each location through the implementation of multiple iterations of filtering.

We first developed a multiscale adaptive thresholding approach inspired by our previous work focused on the sensitive detection limited to the case of diffraction-limited fluorescent structures ^16^. Let us consider the following image model

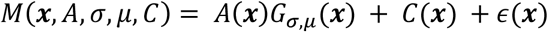

where *A* denotes the spot amplitude, ***x*** the 3D position, *G*_*σ,µ*_(***x***) is a Gaussian function with standard deviation *σ* and mean *µ, C* is the background signal and *ϵ*(***x***) is the additive noise following a Poisson-Gaussian stochastic footprint. The least-square formulation of our optimization problem as

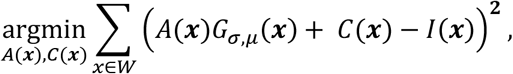

where *I*(·) denotes the image volume and *W* is a 3D box of size 8*σ*, can be simplified to the resolution of a linear system that can be decomposed in multiple filtering passes:

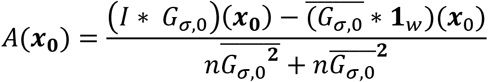

and

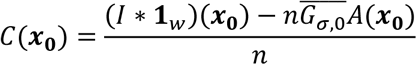

where ***x***_0_ is the fixed voxel position **1**_*w*_ is a unitary convolution kernel along *W*, n is the number of voxel encompassed in *W*. The statistical analysis of the local residuals resulting from the fit

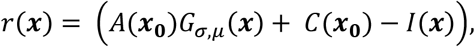

with ***x*** ∈ *W*, provides a p-value-based threshold for testing for the hypothesis that *A*(***x***_**0**_) ≫ *C*(***x***_**0**_) as described in^16^. This approach yields a sensitive binary map *H*_0,*σ*_(·) for the detection of the voxel describing a fluorescence object at scale *σ*. This approach avoids the fitting of an object template in order to reduce computation time.

Next, we carry out this adaptive thresholding step at multiple scale to obtain a vote map.

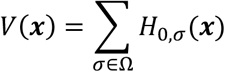

where Ω is the scale range, typically ranging between 120 nm and 1 um. The resulting object mask *V*(·) thus summarizes the presence of particles at any scale at a given voxel (see Supp. Fig. 3) using only filtering operations that can process each voxel in a parallelized fashion. In order to refine the localization of objects present in contiguous object masks, we implemented a multiscale Laplacian of a Gaussian filtering framework^34^ to estimate a map of scale response *S*(·) for each voxel defined as:

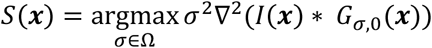

where ∇^2^(·) denotes the Laplacian operator. The watershed algorithm is then applied to further segment this scale response map to detect touching objects. The center of object is determined through the weighted centroid of the voxels belonging to a same object mask.

### Dynamic Region of Interest estimation

In order to visualize and map the molecular processes nested in volumetric time lapse sequences, we propose a framework for the definition of dynamic regions of interest (dynROI) from point cloud sequences. Those dynROIs are described by dynamic bounding boxes (or rectangular cuboids) that are sized to fit the data optimally and oriented according to a moving frame of reference. In this note, we describe the general principles underpinning the estimation of those dynROIs from dynamic point clouds and their implementation across scales: from cellular down to molecular dynROIs.

#### Generic point cloud tracking principle

We first define an optimal frame of reference in the first time point of the sequence with an origin *O*_0_ described by the average point cloud position and with unit vectors (*u*_0_, *v*_0_, *w*_0_) described by the eigenvectors of the covariance matrix of the point positions (a.k.a. principal component analysis). The orientation of the dynROI box in the first frame is described by this frame of reference and its size is defined by the boundaries of the point cloud augmented by a tunable margin (default is set to 5 voxel). The frame of reference at time t is then estimated through a rigid transform as:

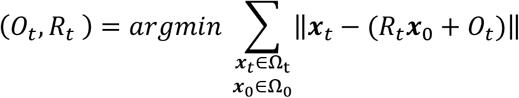

using the Iterative closest point algorithm^35^, where Ω_*t*_ denotes the set of points coordinate at time *t*. The unit vector (*u*_*t*_, *v*_*t*_, *w*_*t*_) are then estimated by applying the rigid transform to (*u*_0_, *v*_0_, *w*_0_). At each time point the size of the box is adjusted to fit the extension of the current point cloud in the current orientation with the additional margin. Multiple dynROI shapes have then been implemented to adjust to the local process (box, sphere, tube, rounded tube, plane and cone).

#### Dynamic region of interest estimation for the cell

The cell is first segmented using the Otsu algorithm and the point cloud representing the cell mask is downsampled randomly to reduce its density by 90% and speed up computations. The generic point cloud tracking principle described above is applied to the downsampled sequence with a margin set to 30 voxels and a box-shaped dynROI (see Supplementary Figure 6).

#### Dynamic region of interest for the spindle

Spindle poles were detected using the multiscale detector with the default p-value (set to 0.005) and scales ranging from 0.4 to 0.8 ums. The motion of poles was modeled with a piecewise stationary Brownian and Directed motion model with a maximum instantaneous displacement set to 3 times the process noise estimated from a Kalman filtering of the trajectory, a lower bound set at 0.5 um and upper bound set at 0.8 um. Failure to detect the very dynamic aggregate on nucleating microtubules is handled with gap closing, the maximum gap is set to 2 s (or 2 frames) with a minimum length of 2 s for the track segment. The resulting dynROI was built has a rounded tube center with a fringe of 9 microns around the segment formed by the two brightest objects present during the complete sequence.

#### Dynamic region of interest for the chromosomes

The kinetochores marking the center of chromosomes were detected using the multiscale detector with the default p-value (set to 0.005) and scales ranging from 0.15 to 0.25 voxels. Motion was modeled with a Brownian motion model with a maximum instantaneous displacement set to 5 times the process noise estimated by Kalman filtering of the trajectory, a lower bound set at 0.4 um and upper bound set at 0.6 um. Variation in SNR were managed with a maximum gap set to 4 s (or frames) with a minimum length of 2 s for the track segment. The dynROI was estimated using the generic point cloud tracking principles described in Section 5.1 using all the trajectories detected inside the spindle dynROI with a box-shaped dynROI and a margin of 0.1 um.

#### Dynamic region of interest estimation for the interpolar region

Let 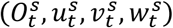 and 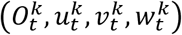 denote the frames of reference estimated for the spindle and the chromosome respectively. We want to build a frame of reference 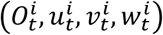 that follows an interpolar plane showing how microtubule nucleation events inside the spindle are orchestrated to capture chromosomes efficiently. We first set the origin to 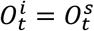 and 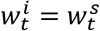 so that one axis is following the spindle at all time. For the plane to describe the motion of the chromosome population, the second unit vector follows a slice of the kinetochore-associated dynROI 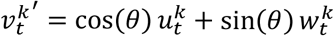 projected to ensure orthogonality as 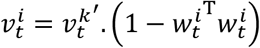. Finally the last unit vector is set as 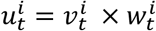. The dynROI type is a plane with a lateral fringe of 50 voxels, a height of 4 voxels and an angle *θ* set to 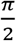.

#### Dynamic region of interest estimation for the Kinetochore fibers

Assuming K-fibers to span the region between poles and kinetochores as a straight polymer, its associated microtubule dynamics was observed using a conical dynROIs with an angle of pi/12.

### Stochastic programming for the evaluation of trackability

The association of particle detections with trajectory heads is performed in a temporally greedy fashion, i.e. particles detected at time *t* are linked to the heads of track segments defined up to time *t* − 1 without consideration of the track segments beyond *t* and only indirect consideration of track segment before t-1. Therefore, our definition of trackability relates to the level of ambiguity in assigning particles detected in time point *t* to track segment heads in *t* – 1. The optimal association is obtained by linear assignment of heads to particles in a bi-partite graph:

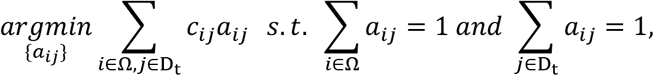

where Ω is the set of track segment heads, *D*_*t*_ is the set of detections measured at time *t, a*_*ij*_ ∈ {0,1} denotes the assignement of the *i*th track segment to the *j*th particle and *c*_*ij*_ ∈ ℝ is the cost associated to making that association. The association cost *c*_*ij*_ typically reflects the distance between the predicted location of the *i*th track segment at *t* and the *j*th detection at this same time point. This assignment problem is convex, hence with a guaranteed unique solution, and can be solved using a variety of linear programming algorithms^15,36,37^. However, a key challenge in our framework is the deterministic aspect of this solution. There is no measure of uncertainty attributed to the final graph of associations (see Fig 5.a). While several algorithms have been proposed to estimate the uncertainty related to the total optimal cost of a linear programming problem, a.k.a. stochastic programing ^38^, they do not focus on the detection of local changes in association made in the bi-partite graph. In this Section, we will first detail how we consider the randomness present in the history of each track to estimate the probability distribution associated to all assignment costs *c*_*ij*_. We will then describe how these uncertainties can then be exploited to detect local ambiguities in the assignment problem, which subsequently define a score of trackability.

Stochastic filtering approaches are routinely used to estimate the parameter describing the dynamic properties of tracked particles from their position history. They enable the prediction of particle location from one frame to the next to refine the cost used for linear assignment. Those temporally recursive algorithms also provide inferences of track segment prediction uncertainty from *t*-1 to *t*. Briefly, let ***x***_*t*_ be a variable describing the state of the track segment. For a particle moving in a directed fashion, it is defined as:

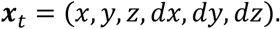

The associated probability *p*(***x***_*t*_ |**z**_1:*t*_) can be estimated recursively thanks to the Bayes rule:

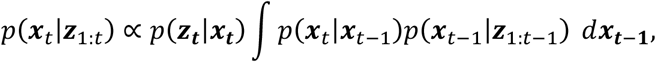

where **z**_**1**:***t***_ represents the past measured positions assigned to a particular track. Kalman filtering is a scalable and flexible way to model the motions of thousands of particles in parallel, and as such is used in the majority of tracking approaches^7^, including u-track. In this framework, the relationships between random variables are assumed to be linear and described as follows:

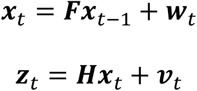

where ***F*** is the state transition matrix between consecutive time points, ***H*** is the observation matrix, and ***w***_*t*_ and ***v***_*t*_ are the model and measurement noise respectively, both assumed to be Gaussian with covariance matrices ***Q***_*t*_ and ***R***_*t*_. The Gaussian and linear assumption provides an analytical solution with a computationally efficient implementation to estimate 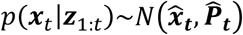 (see our previous work^14^ for a detailed review). Before optimal assignment between a track segment at *t* − 1 and the object detected on frame *t*, the probability distribution of the predicted particle positon at time *t* is then described by 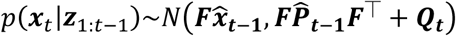. As such the variation of the cost to associate the *i*th track segment to the *j*th measurement can be expressed, without loss of generality as:

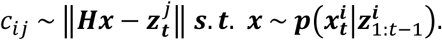

This expression provides us with a direct way to explore the space of possible combination of cost values through Monte Carlo Simulations. U-track 3D implements several types of stochastic filtering approaches such as unimodal and multimodal Kalman filtering as well as piecewise stationary motion filtering or smoothing approaches, where the same principles can be straightforwardly applied. The principle underlying the use of our predicted probability distribution to evaluate assignment stability is described graphically in Figure 5. Our local trackability score is defined as:

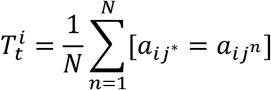

where *a*_*ij*_* is the initial assignment found for the *i*th trajectory, *a*_*ij*_*n* is a newly computed assignment resulting from the *n*th out of a total of *N* simultaneous resampling rounds of all costs *c*_*ij*_ and [.] denotes the Iverson bracket. Each new assignment result, or vote, is considered different if the track segment is assigned to another detection, or determined to be a track termination. As such, a lower score 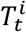 reflect a larger instability in the optimal assignment, hence a higher ambiguity and lower trackability. In our experiments, the number of resampling rounds is set to *N* = 20.

### Simulation parameters for trackability evaluation

Parameters used to simulate the object described in Section “The trackability score detects tracking ambiguities with high precision”.

| Varying parameter | Diffusion | Density | Velocity | Transition Rate |
| --- | --- | --- | --- | --- |
| Figure | 5.f-i | Supp 2 | Supp 3 | Supp 4 |
| Diffusion Coef. ($\mu\text{m}^2/\text{s}$) | 0.1 – 1.0 | 0.3 | 0.15 | 0.1 |
| Density ($\mu\text{m}^{-3}$) | 0.1 | 0.01 – 0.5 | 0.1 | 0.2 |
| Velocity ($\mu\text{m}/\text{s}$) | 0 | 0 | 0 – 2.2 | 1.5 |
| Transition Rate | 0 | 0 | 0 | 0 – 0.5 |
| Volume size ($\mu\text{m}^3$) | 1000 | 1000 $\mu\text{m}^3$ | 1000 $\mu\text{m}^3$ | 1000 $\mu\text{m}^3$ |
| Frame rate (Hz) | 1 | 1 | 1 | 1 |
| Movie Length (s) | 150 | 150 | 150 | 150 |
| Trajectory lifetime (s) | 20 | 20 | 20 | 20 |
| Standard error on detection | 0.005 | 0.005 | 0.005 | 0.005 |

### Clathrin-mediated endocytosis study on a glass coverslip

#### Cell preparation and imaging

Inner medulla collecting duct (IMCD) mouse epithelial cells (ATCC CRL-2123) stably expressing alpha-adaptin GFP^16^ were cultured in DMEM/F12 supplemented with 10% fetal calf serum and 1% antibiotic/antimycotic. Cells were plated on 5 mm diameter coverslips (64-0700, Thomas Scientific) and mounted to a custom machined holder for imaging with a high-NA version of diagonally scanned light-sheet microscopy^2^. This microscope is equipped with an NA 0.71 water dipping illumination objective (54-10-7, Special Optics), and a 25X/NA 1.1 water dipping detection objective (CFI75 Apo LWD 25XY, Nikon Instruments), and a Hamamatsu Flash 4.0 sCMOS camera. Briefly, 500 time points were acquired with 30 uW of 488 nm illumination (measured at the back pupil of the illumination objective) and a 15 ms camera exposure. Each image stack was 106.5 × 39.9 × 23.1 um^3^, with a lateral and axial voxel size of 104 and 350 nm, respectively, resulting in a 1.008 Hz volumetric image acquisition rate.

#### Clathrin structure trajectory estimation and post-processing

Clathrin structure aggregates, labelled by alpha-adaptin GFP were detected using a multiscale particle detector with a p-value set to 0.05 and scales ranging from 0.15 um to 0.5 microns. For tracking, the motion of particles was modeled with a Brownian motion model with a maximum instantaneous displacement set as three times the process noise estimated by Kalman filtering of the trajectory, a lower bound set at 0.1 um and upper bound set at 0.3 um. When detection gaps are enabled, the maximum gap length is set to 3 s (or frames) with a minimum length of 3 s for any track segment allowable to be connected by the gap closing algorithm^39^.

The median of the maximum intensity reached per track was then used to discriminate between abortive and maturing CCPs. To account for the variation of fluorescence signal across acquisitions, the maximum intensities were scaled such that the empirical cumulative distribution function (cdf) of maximum intensities computed for each acquisition matched the median cdf of all acquisitions, as previously described in ref^16^.

### Microtubule instability measurement

#### Cell preparation and imaging

Cell preparation and imaging of microtubule plus-ends has been carried with Lattice Light-sheet microscopy as described in ref^1^.

#### Plus-ends trajectory estimation

Plus-ends, labelled through GFP tagging of EB1, were detected using a multiscale detector with the default p-value (set to 0.005) and scales ranging from 0.15 um to 0.25 um. The polymerization of microtubule was modeled with a directed displacement estimated through a Kalman filtering of the trajectory, similar to^18^, but now in 3D. The random component of this displacement was estimated as 3 times the process noise of the Kalman filter with a lower bound of 0.3 um and an upper bound of 0.6 um.

The shrinkages and pauses detection framework proposed in^18^ has also been translated to 3D plus-ends trajectories. The detection of both shrinkages and pauses is carried out by closing gaps between track segments, which implicitly delineate phases of microtubule growth. In our experiment, the minimum growth duration to consider gap closing was set to 4 s (or frames) and the maximum gap duration was set to 8 s. In order to detect pauses, the maximum angle between a speed vector estimated immediately prior and posterior to the pause event was set to 30 degrees, the maximum positional fluctuation in the plus-ends location during a pause is set to 0.5 um. To detect a shrinkage event between two segments, we first measure the distance D between the termination point of the earlier segment (which is equivalent to the potential locus of a catastrophe event) and the initiation point of the later segment (which is equivalent to the potential locus of a rescue point) along the path of the earlier segment. The two segments are connected by the gap closer if the distance between the initiation point to the closest point along the trajectory of the first segment does not exceeds *Dsin*(*θ*) with *θ* set to 20 degrees in our experiment.

### Single molecule dynamics study with lattice light-sheet microscopy

#### Cell preparation and imaging

Cell preparation of fluorescently labelled Sox2 transcription factor for single molecule imaging is described in ref^1^. Briefly, 100 time points were acquired with Lattice light-sheet microscopy imaging. Nine planes spaced 500nm apart were acquired at 50 ms of camera exposure, resulting in a 2 Hz volumetric image acquisition rate. Each image stack was 50 × 50 × 5 um^3^, then cropped around the nucleus, with a lateral and axial voxel size of 100 and 500 nm, respectively.

#### Estimation of transcription factor binding times

Transcription factor single molecules were detected using a multiscale detector using a p-value of 0.01 and a scale ranging from 0.15 to 0.3 um. Transcription factor motion was modeled using a Brownian motion model with a maximum instantaneous displacement estimated as 6 times the process noise estimated by Kalman filtering of trajectory to account for speed variations during long periods of confined diffusion, a lower bound set at 0.3 um and upper bound set at 0.5 um. The maximum gap is set to 4 s (or frames) with a minimum length of 2 s for any track segment allowable to be connected by the gap closing algorithm. We assume that if the single molecule is detectable it immobilized at the DNA. Accordingly, characteristic binding times *τ* are estimated by a double exponential fit to the lifetime distribution.

### Adhesions and collagen interaction imaging and analysis

#### Cell preparation and imaging

For three-dimensional imaging of adhesions, mycoplasma-free U2OS cells were cultured in DMEM with 10% fetal bovine serum (Sigma; F0926-500ML) at 5% CO2 and 37 °C. Cells were lentivirally transduced with a truncated CMV promoter (Addgene #110718) driving the expression of mNeonGreen-Paxillin (Allele Biotechnology). Cells were seeded into a pH-neutralized collagen solution (∼2 mg/mL) that, when polymerized, fully embedded cells in a three-dimensional extracellular matrix environment. For visualization of the extracellular matrix, a small concentration of the collagen was fluorescently conjugated with Alexa Fluor 568 NHS Ester (A20003, ThermoFisher). Samples were imaged with a high-NA variant of Axially Swept Light-Sheet Microscopy using 488 nm and 561 nm lasers for illumination (OBIS LX, Coherent, Inc.). The details of this microscope will be published elsewhere. Briefly, lasers are combined, spatially filtered, expanded, and shaped into a light-sheet with a cylindrical lens. This light-sheet was relayed to a bidirectional scan unit (6215, Cambridge Technology), a remote focusing system (CFI S Plan Fluor ELWD, Nikon Instruments), and eventually to the illumination objective (54-10-7, Special Optics). Fluorescence was detected in a widefield format with a water-dipping objective (CFI75 Apo LWD 25SW, Nikon Instruments) and imaged onto two sCMOS cameras (ORCA-Flash4.0, Hamamatsu Photonics) with a 500 mm achromatic doublet (49-396, Edmund Optics), laser line filter, a dichroic, and bandpass filters (ZET405/488/561/640, ZT568rdc, ET525/50m, and ET600/50m, Chroma Technology Corporation). The laser laterally dithered for shadow reduction and scanned synchronously with the detection objective (P-603.1S2 and E-709.SRG, Physik Instrumente) to acquire a three-dimensional stack of images. All equipment was controlled with custom LabVIEW software, which is available from UTSW upon completion of a material transfer agreement.

#### Adhesion detection and elongation analysis

Paxilin aggregates as a surrogate for adhesions were detected using the multiscale detector described in Section “Multiscale particle detector” based on a p-value of 0.001 and a scale ranging from 0.3 to 0.5 um. The elongation of each detected adhesion is computed through a tubularity metric evaluated for each voxel and averaged across all the voxels associated to a single adhesion. Similar to the classic vesselness estimator by Frangi and colleagues^40^, our tubularity metric is based on the eigen values of the

Hessian matrix to describe local curvature,. Let (*λ*_1_ < *λ*_2_ < *λ*_3_) be the three eigenvalues of the Hessian matrix computed at each voxel, the tubularity metric *T* = 1 − | *λ*_1_/*λ*_2_| yields a value between 0 and 1 increasing with the elongation of the adhesions. As such, a noteworthy difference between the classic score described in Frangi’s approach is the use of the two lowest eigen-values (associated with the two axis of lowest curvature direction) to discriminate between flat and elongated adhesions.

### Endosome trackability on cell cultured on top of collagen

#### Cell preparation and imaging

Sum159O breast cancer cells^41^ stably expressing alpha-adaptin GFP were imaged similarly to the one platted on glass coverslip, with the exception that they were plated on a ∼2 mm thick bed of rat tail-derived Collagen Type I (354236, Corning).

#### Clathrin structure trajectory estimation

Clathrin structure aggregates, labelled by alpha-adaptin GFP were detected using a multiscale particle detector with a p-value set to 0.01 and scales ranging from 0.125 to 0.5 microns. For tracking, the motion of particles was modeled with a Brownian motion model. In order to follow the erratic displacements caused by large protrusive motions, the maximum instantaneous displacement was set to 5 times the process noise estimated by Kalman filtering of the trajectory, a lower bound set at 0.3 um and upper bound set at 0.6 um. The maximum gap length is set to 3 s (or frames) with a minimum length of 3 s for any track segment allowable to be connected by the gap closing algorithm^39^.

## Movies Descriptions

### Movie 1

Detail of a rat kidney cells layer expressing GFP-AP2 subunit imaged with diagonally scanned light-sheet microscopy (diaSLM) acquired with a volumetric frequency of 1 Hz and rendered with maximum intensity projection (MIP). Overlay highlights trajectories colored according to maturing events (green), aborting events (orange) and tracks discarded because their lifetimes are cut by the acquisition beginning and end (blue). The red spheres highlight gap location.

### Movie 2

Details of HeLa cell in interphase expressing GFP-EB1 imaged with lattice light-sheet microscopy, rendered with MIP, and acquired with a volumetric frequency of 1 Hz under control condition (left) and after treatment with 33 nM of nocodazole (right). Overlay highlights trajectories, colored uniquely, and yellow circles highlight location of pauses and rescues.

### Movie 3

HeLa cell in metaphase expressing GFP-EB1 imaged with lattice light-sheet microscopy, rendered with MIP, and acquired with a volumetric frequency of 1 Hz. Overlay highlights trajectories. Rendering using the Amira rendering software.

### Movie 4

Dual-colored orthogonal MIP of osteocarcinoma cells expressing eGFP-labeled paxillin and embedded in collagen labelled with Alexa fluor 568 acquired with a volumetric frequency of 0.07 Hz. Overlay highlights dynamic region of interest (dynROI).

### Movie 5

Mask of detected adhesions colored according to their proximity to the closest collagen fiber.

### Movie 6

Mask of detected adhesions colored according to their elongations.

### Movie 7

Slice of the mask of detected adhesions and collagen fibers taken at the center of cell dynROI described in Fig. 2.a and Movie 3.

### Movie 8

Dual-colored orthogonal MIP of HeLa cells undergoing mitosis labeled with eGFP-labeled EB3 and mCherry-labeled CENPA acquired with a volumetric frequency of 0.1 Hz. From left to right, overlays highlight a dynROI built around centrosome trajectories, a dynROI built around CENPA trajectories, and a plane built to visualize the dynamics of chromosomes relative to the spindle location.

### Movie 9

Dual-colored orthogonal MIP of HeLa cells undergoing mitosis labeled with eGFP-labeled EB3 and mCherry-labeled CENPA acquired with a volumetric frequency of 0.1 Hz. The red overlay highlights the dynROI tracking a plane built to visualize the dynamics of chromosomes relative to the spindle location.

### Movie 10

Left: Dual-colored orthogonal MIP of HeLa cells during pro-metaphase acquired with a volumetric frequency of 1 Hz. Overlay highlights the motion of the dynROI. Right: Rendering of the volume described by the dynROI in its associated frame of reference. Green dots highlight plus-ends inside the mapping cone and red circle describe the motion of the target kinetochore.

### Movie 11

Top: Orthogonal MIP of breast cancer cells imaged with diaSLM expressing eGFP-labelled alpha subunit of the AP-2 complex acquired with a volumetric frequency of 1 Hz. Rectangular overlays show dynROIs with different types of dynamic of activity. Dot overlays show local level of ambiguity. Bottom: Rendering of the volume described by the dynROI in their associated frames of reference.

---

**Supplementary Figure 1:**
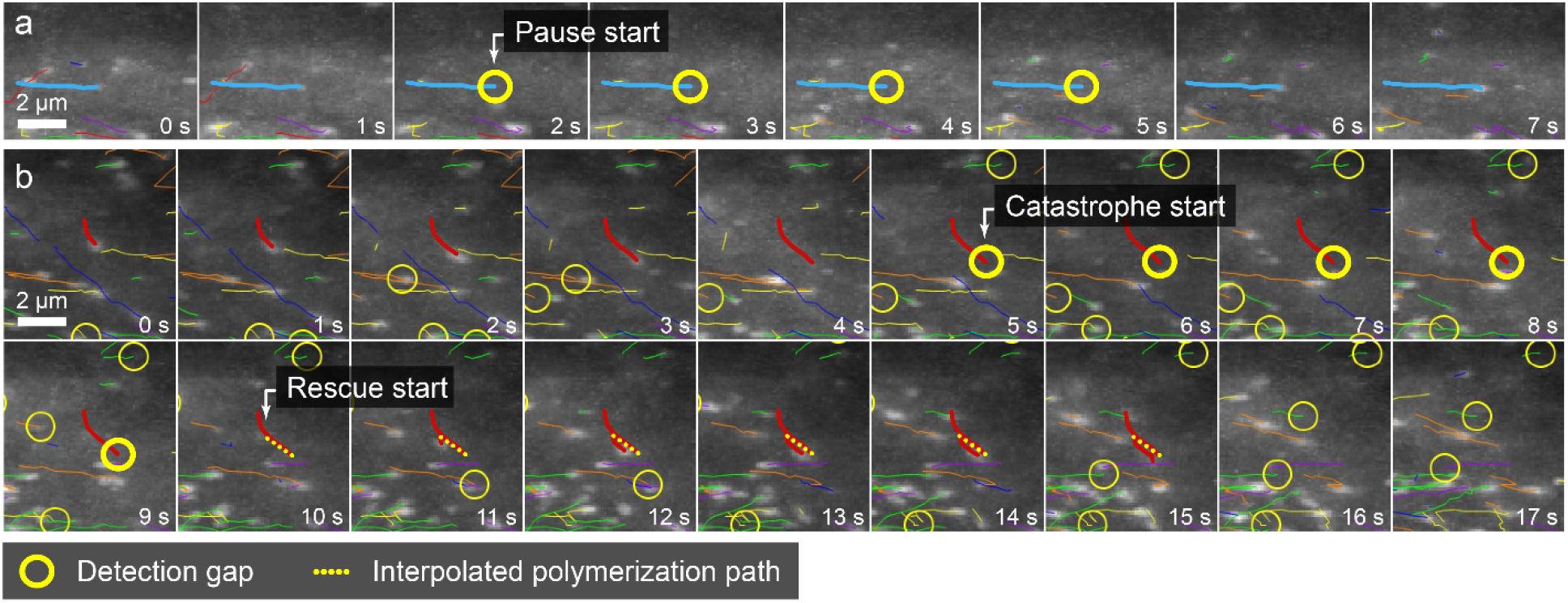
a) Example of a pause in microtubule polymerization detected in HeLa cell in interphase (detail). Yellow circles highlight detection gaps. b) Example of catastrophe and rescue events detected in the same sequence (detail).

**Supplementary Figure 2:**
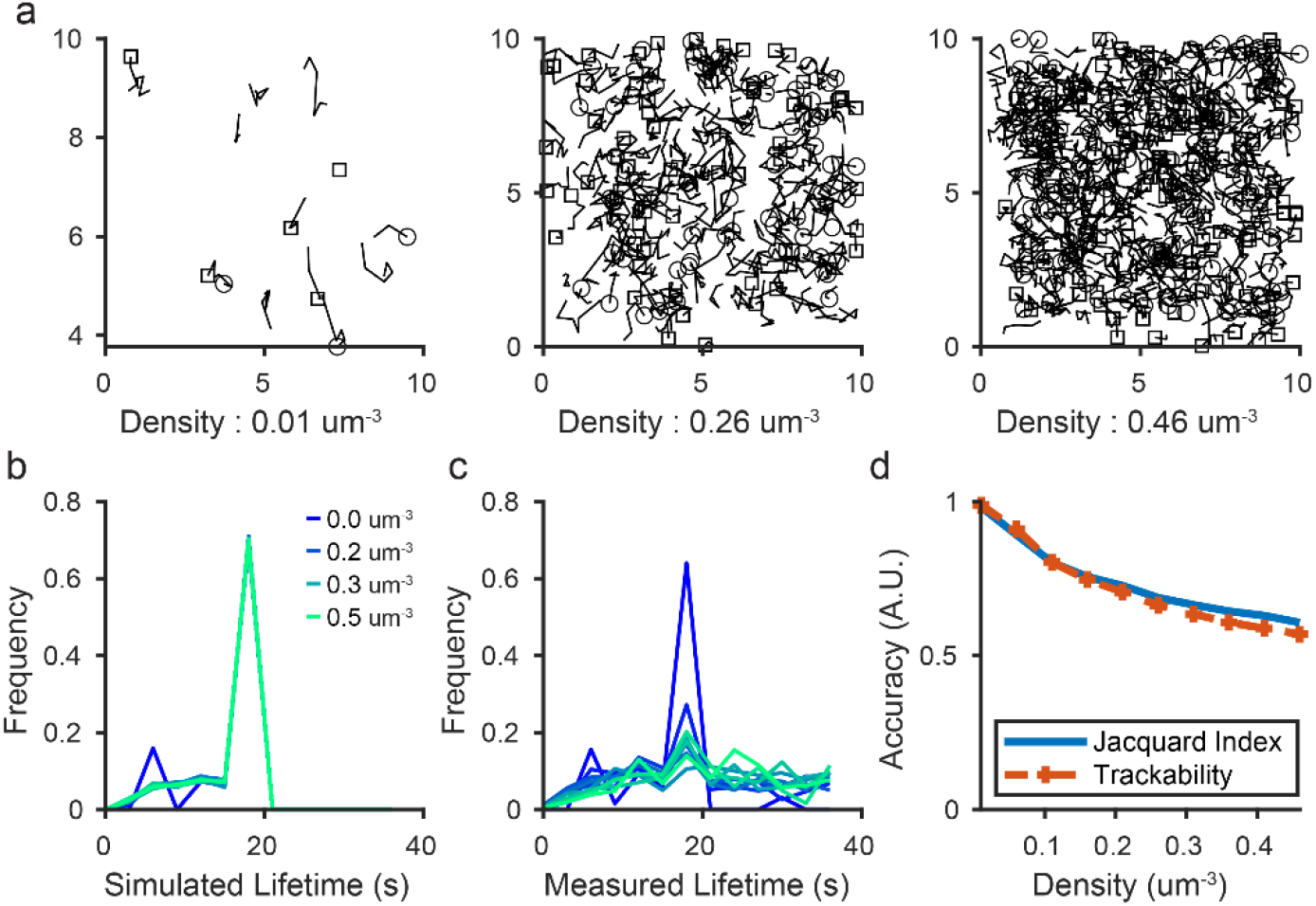
The trackability score predicts the performance decrease associated to particle density. a) Examples of simulated trajectories with particle density ranging from 0.01 to 0.5 um^-3^ with a fixed diffusion coefficient of 0.3 um^2^/s. Visualization is limited to five consecutive frames to reduce clutter. b) Lifetime of simulated trajectories. c) Lifetime distribution measured through tracking. d) Accuracy measured through the jacquard coefficient on the ground truth and estimated with the trackability score using the detection set.

**Supplementary Figure 3:**
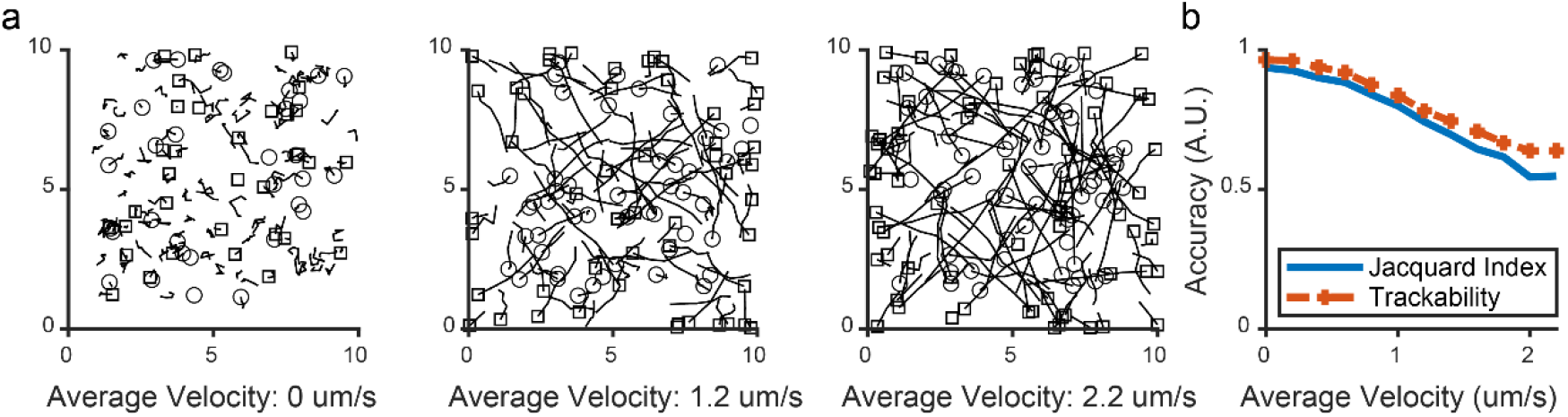
The trackability score predicts the decrease in performance associated to particle velocity. a) Examples of simulated trajectories presenting directed motions described by velocities ranging from 0 to 2.2 um/s with a fixed diffusion component coefficient of 0.15 um^2^/s and density set to 0.1 um^-3^. Visualization is limited to five consecutive frames to reduce clutter. b) Accuracy measured through the jacquard coefficient on the ground truth and estimated with the trackability score using the detection set.

**Supplementary Figure 4:**
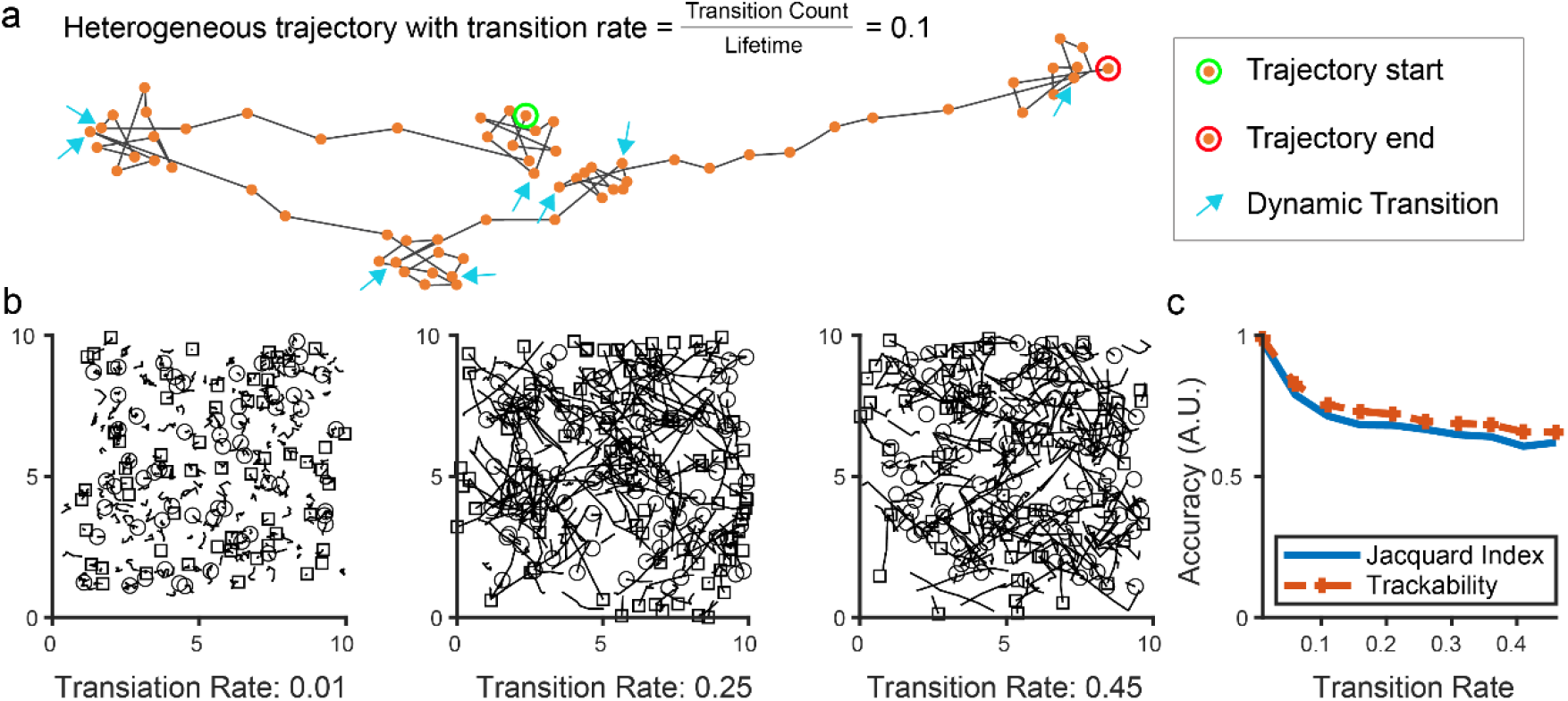
The trackability score predicts the decrease in performance associated to the heterogeneity of motion types in a single trajectory. a) Illustration of the transition rate used to simulate a dataset with increasing heterogeneity. b) Examples of simulated trajectories with diffusive motion described by a coefficient set to 0.1 um^2^/s, and directed motion set to 1.5 um/s with a diffusive component of 0.1 um^2^/s. Density is set to 0.2 um^-3^. Visualization is limited to five consecutive frames to reduce clutter. c) Accuracy measured through the jacquard coefficient on the ground truth and estimated with the trackability score using the detection set.

**Supplementary Figure 5:**
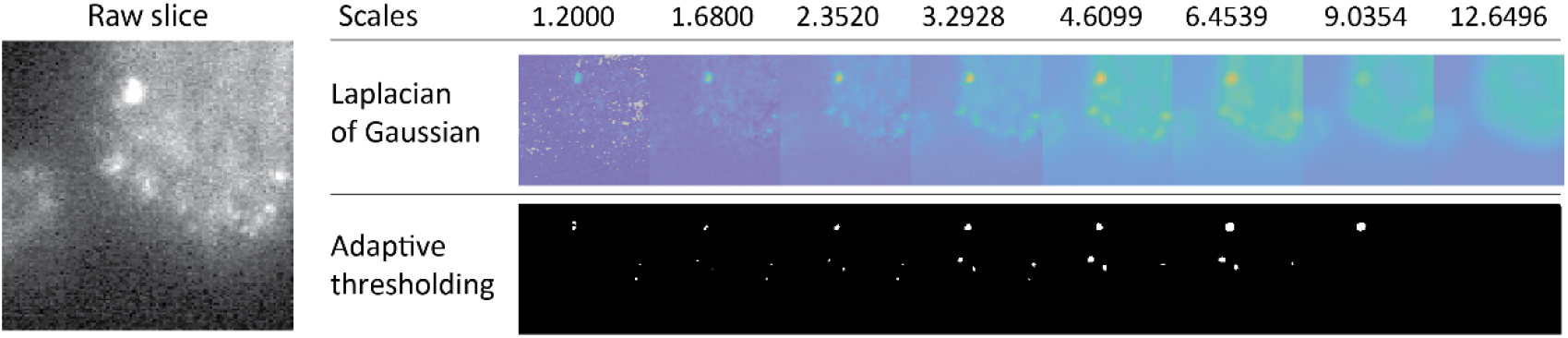
Principle of multiscale Laplacian-of-Gaussian filtering (top) and multiscale adaptive thresholding approach (bottom) demonstrated on a slice of a volumetric imaging of cellular adhesions (detail).

**Supplementary Figure 6:**
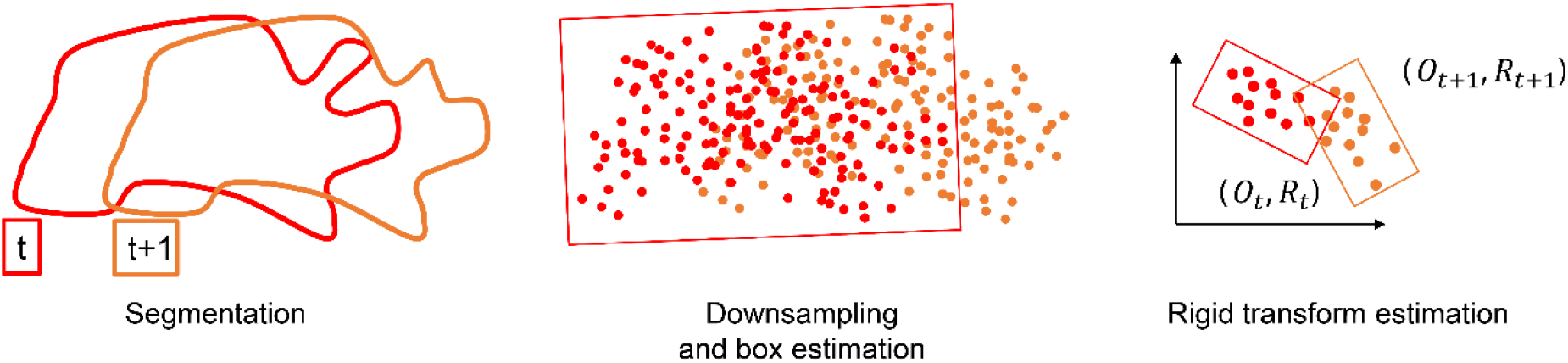
Principle of generic point cloud tracking applied to cell Dynamic region of interest estimation for the cell.

## Notes

https://github.com/DanuserLab/u-track3D

### Competing Interest Statement

The authors have declared no competing interest.

### Summary of Updates

The evaluation of the trackability score has been significantly improved. We now show the high predictive power on tracking error using simulations. We have also split figures to improve their focus on a single message.

